# BamHash: a checksum program for verifying the integrity of sequence data

**DOI:** 10.1101/015867

**Authors:** Arna Óskarsdóttir, Gísli Másson, Páll Melsted

## Abstract

**Summary:** Large resequencing projects require a significant amount of storage for raw sequences, as well as alignment files. Since the raw sequences are redundant once the alignment has been generated, it is possible to keep only the alignment files. We present BamHash, a checksum based method to ensure that the read pairs in FASTQ files match exactly the read pairs stored in BAM files, regardless of the ordering of reads. BamHash can be used to verify the integrity of the files stored and discover any discrepancies. Thus, BamHash can be used to determine if it is safe to delete the FASTQ files storing raw sequencing reads after alignment, without the loss of data.

**Availability and Implementation:** The software is implemented in C++, GPL licensed and available at https://github.com/DecodeGenetics/BamHash

## 1 INTRODUCTION

Resequencing projects, where individuals are sequenced from a species with a known reference genome, generate a significant amount of raw sequences that are then aligned to the reference genome. Data storage becomes an issue as the cost of sequencing decreases and the throughput of current sequencing technologies keeps increasing.

Raw sequencing reads are generally stored in FASTQ file format, usually compressed. After read mapping the resulting alignment is stored in a BAM file (Li *et al.,* 2009). This BAM file is then sorted and processed further, but most importantly it contains all the original information of the FASTQ file. Sorted BAM files yield a better compression, compared to unsorted BAM files, as well as allowing random lookup over genomic regions. For this reason almost all post-alignment analysis, e.g. variant calling, realignment, and local assembly are done on the sorted BAM file, rather than the original FASTQ file.

Since the BAM file contains all the information of the FASTQ file it is justifiable to delete the FASTQ file after alignment. After all, the contents of the FASTQ file can be regenerated from BAM file.

However, before deleting the FASTQ file, we need to be sure that there is no loss of data, i.e. that the sequences in the FASTQ file are exactly the same as the sequences in the BAM file. The two files could differ due to a number of reasons. Any errors in the alignment pipeline could generate inconsistent files.

We present BamHash, a tool for verifying the data integrity between a FASTQ and a BAM file. The program computes a 64-bit fingerprint from the sequences and read names for both FASTQ and BAM files. The method is highly sensitive to changes in the input so a change in a single nucleotide will result in different fingerprints; the probability of generating the same fingerprint by chance is astronomically small. The role of this tool is to flag any FASTQ and BAM files that have different fingerprints and mark the FASTQ files as unsafe for deletion.

BamHash plays the same role as the md5sum program, which computes a fingerprint of files. Comparing md5sum fingerprints (Rivest, 1992) of FASTQ and BAM files would not yield a comparable result, since the formatting and ordering are different. Our method is fast and memory efficient; it can compute the fingerprint of a BAM file from 30-fold coverage human sequencing experiment in 30 minutes.

## 2 METHODS

The information for the sequencing reads, namely the read name, sequence and quality values are stored differently in FASTQ and BAM files, but can easily be parsed and recovered. The internal order of reads is generally not conserved, unless guaranteed by the alignment software. Since BAM files sorted by genomic coordinates are the norm, we cannot expect to maintain the order.

Thus we need to compare the two files as sets, or rather multisets, of items. To do this we use a hash function *h* for each item and reduce all hash values using a commutative binary operation. The commutative property ensures that the final result is independent of the ordering of the reads.

For the commutative binary operation, the XOR function is a natural candidate for sets. However, XOR has the property that each value is it’s own inverse, if an item *x* is repeated twice in the input, the hash values will cancel each other since *h*(*x*) ⊕ *h*(*x*) = 0. Normally, we do not expect to see repeated items since read names tend to be unique, however if the read names have been stripped and quality values are absent we cannot guarantee that this holds. For this reason we chose to work with the sum of hash values as 64-bit integers.

By using a hash function we ensure that the resulting fingerprint is sensitive to any changes in the input. For BamHash, we chose the MD5 hash function. Whereas MD5 was developed for cryptographic purposes, making it hard to forge an MD5 signature, we only rely on the sensitivity of the hash function to catch accidental changes. It should be noted that the proposed method cannot guarantee that it would be too hard for a malicious agent to modify the input to produce any given fingerprint.

### 2.1 Implementation

The pseudo-code for the method is given in Algorithm 1. For FASTQ files, the input for paired reads is given by two FASTQ files. Each read in the FASTQ file is processed and the read names for pairs is modified to end in /1 or /2, if the read wasn’t already in this format. For BAM files, a single BAM file is given and the flags are used to determine whether each read is the first or second in a read pair. Furthermore, reads that are mapped to the reverse strand have the reverse complement of the sequence stored in the BAM file to aid compression. For these reads we reconstruct the original sequence to match what was stored in the corresponding FASTQ file. The program is written in C_++_ and uses the SeqAn library (Döring *et al.,* 2008) for parsing FASTQ, gzip compressed FASTQ and BAM files.

#### Algorithm 1 Checksum for paired sequences

 **function** Hash-update-Bam(*r*)

          *s* ← *r*.name

          **if** *r* is first in pair **then**

                  *s* ← *s*+ “/1”

          **else**

                  *s* ← *s*+ “/2”

          **if** *r* is on reverse strand **then**

                  *r*.seq ← REVERSE-COMPLEMENT(*r*.seq)

          *s* ← *s* + *r*.seq + *r*.qual

          **return** MD5(*s*)

 

**function** Hash-update-Fastq(*r*)

          *s* ← r.name + *r*.seq + *r*.qual

          **return** MD5(*s*)

 

**function** Hash-file(*f*)

          *H* ← 0

          **for all** reads *r* in *f* **do**

                  **if** *f* is a BAM file **then**

                         *H* ← *H* + (Hash-update-Bam(*r*)) mod 2^64^

                  **else**

                         *H* ← *H* + (Hash-update-Fastq(*r*)) mod 2^64^

          **return** *H*

## 3 RESULTS

To assess the performance of BamHash, we compared the running time for processing BAM files to viewing with Samtools. The dataset chosen was a whole genome sequencing experiment, aligned to GRCh38 Human reference using BWA-MEM. All datasets were generated at the laboratory at deCODE Genetics and were processed with the same pipeline (Gudbjartsson *et al.,* 2014). The BAM file consists of 832 million read pairs at 38× coverage. BamHash required 38 minutes to compute the hash values, whereas Samtools required 49 minutes to parse the BAM file and count lines. We note that the program is largely I/O bound. It runs on a single core, which is underutilized, as most of the time is spent waiting for data from the disk.

## 4 DISCUSSION

The role of BamHash is to detect differences between the read sets of raw FASTQ and aligned BAM files. This discrepancy can arise due to mistakes in the pipeline, bugs in alignment code or disk failures. When the data integrity has been verified, the original FASTQ files can be safely discarded, thus freeing up storage space. Additionally, BamHash will be useful when porting alignments to a new reference genome. Such a pipeline would create intermediate FASTQ files, which would then be aligned to the new reference. The old BAM file can be removed only if the BamHash signature agrees with the newly created alignment.

## REFERENCES

Döring, Andreas, David Weese, Tobias Rausch, and Knut Reinert. SeqAn an efficient, generic C++ library for sequence analysis. BMC bioinformatics 9.1 (2008): 11.

Daniel F. Gudbjartsson et al. Large-scale whole-genome sequencing of the Icelandic population. Nature Genetics. In press.

Li, Heng, Bob Handsaker, Alec Wysoker, Tim Fennell, Jue Ruan, Nils Homer, Gabor Marth, Goncalo Abecasis, and Richard Durbin. The sequence alignment/map format and SAMtools. Bioinformatics 25.16 (2009): 2078–2079.

Rivest, Ronald. RFC 1321: the MD5 message-digest algorithm. Internet Engineering Task Force (1992).

